## supplementary materials for "Predictive evolutionary modelling for influenza virus by site-based dynamics of mutations"

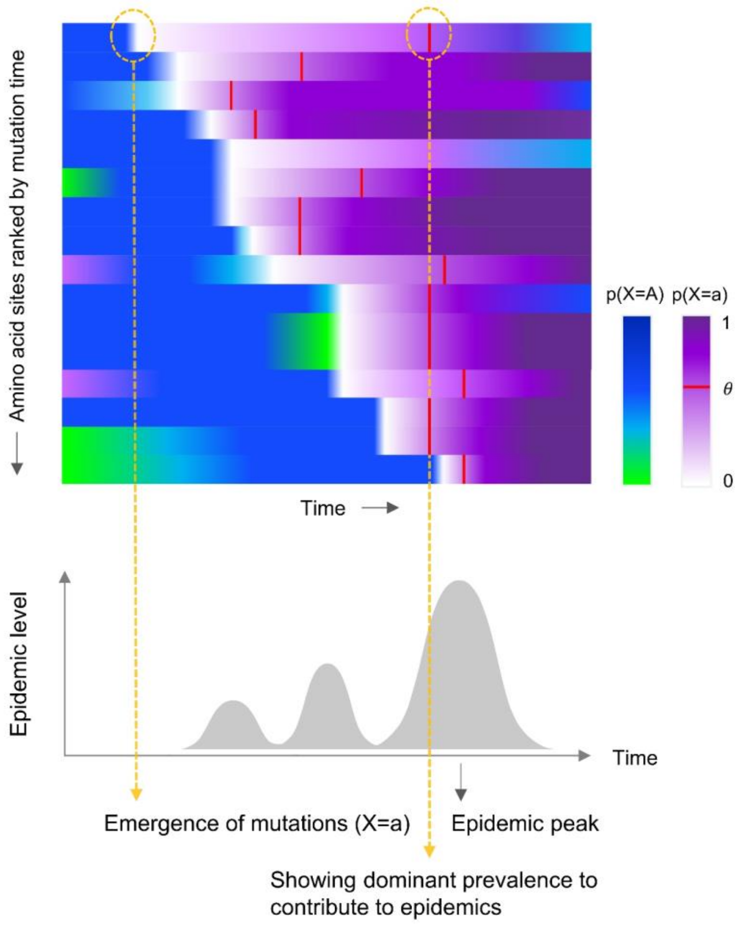


**Extended Data Fig. 1 | Schematic plot of the relationship between mutation prevalence dynamics in genome and population epidemics.** Each color represents the dynamic change of a mutations at a genomic site (by row). Mutations with selective advantage emerge and lead to viral fitness at varying paces. For example, in the first row of the upper panel, P($X$ =$a$), the mutation prevalence of a residue $a$ at a codon position $X$ gradually increases from 0 (white) to 1 (purple). $\theta$ is the mutation prevalence threshold indicating that the substitution is detected to be significantly associated with the rise of epidemics level. P($X$ =$A$) is the prevalence of prior residue $A$ at position $X$, ranges from 0 (green) to 1 (blue). The proposed method draws information from the mutation transition time to model the dynamics of evolution.

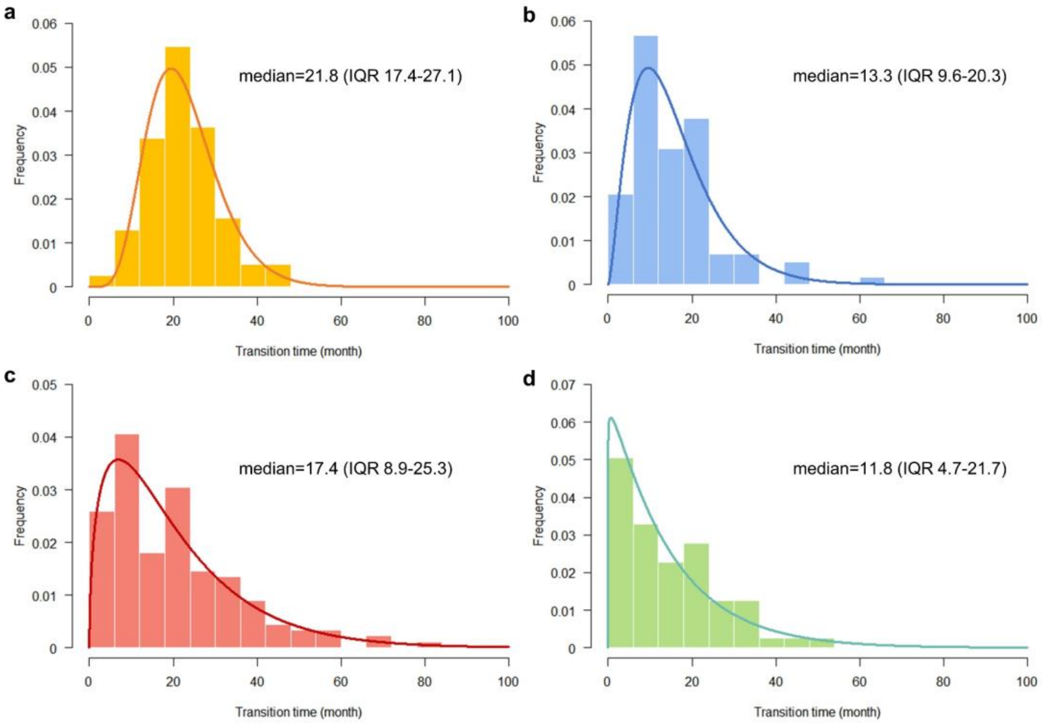


**Extended Data Fig. 2 | Empirical probability distribution of transition time $\boldsymbol{\tau}$.** a, pH1N1 HA. b, pH1N1 NA. c, H3N2, HA. d, H3N2 NA. The transition time $\boldsymbol{\tau}$ of mutation from emergence to an influential threshold θ for population epidemics exhibits gamma-like distribution.

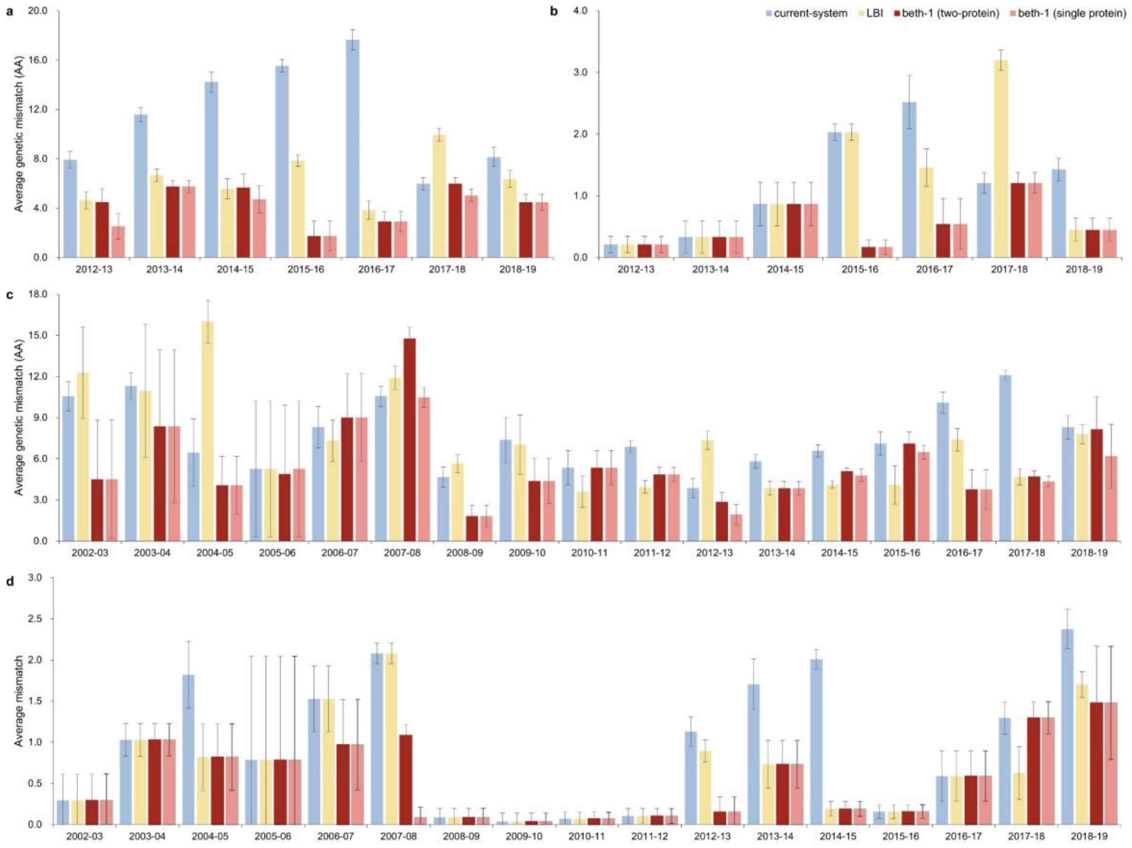


**Extended Data Fig. 3 | Prediction accuracy of alternative methods for NA evolution in the retrospective data by season.**

**a**, pH1N1, NA full sequence (469 codons). **b**, pH1N1, NA epitopes (30 sites). **c**, H3N2, NA full sequence (469 codons). **d**, H3N2, NA epitopes (12 sites). Prediction accuracy is assessed by the average amino acids mismatch (Y-axis) of the predicted strains against circulating viruses in ten geographical regions in the respective epidemic seasons (X-axis). Error bar: standard deviation of regional mean mismatch.

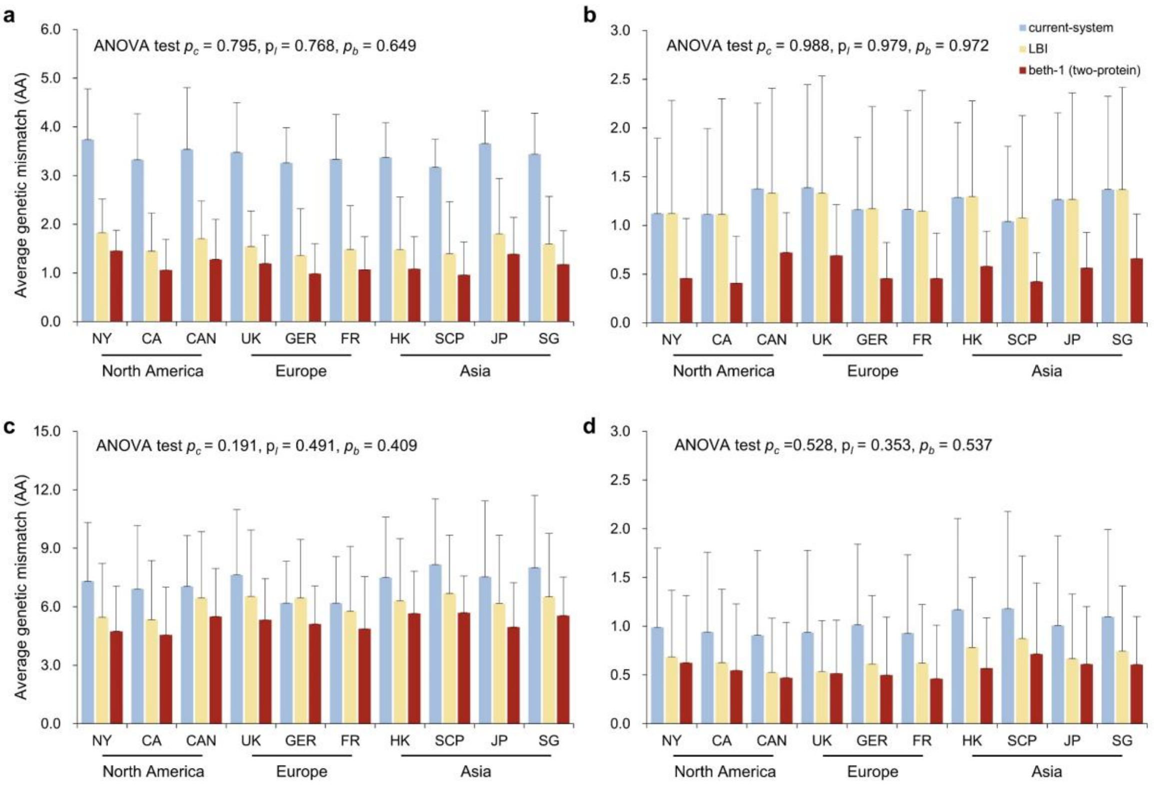


**Extended Data Fig. 4 | Prediction accuracy on epitopes, breakdown by geographical region. a**, pH1N1, HA epitopes. **b**, pH1N1, NA epitopes. **c**, H3N2, HA epitopes. **d**, H3N2 NA epitopes. $p_c$: $p$-value of ANOVA test on equality of genetic mismatch across three continents by the current-system; $p_b$: by beth-1 (two-protein). No significant difference in genetic mismatch are observed across continents using either prediction models.

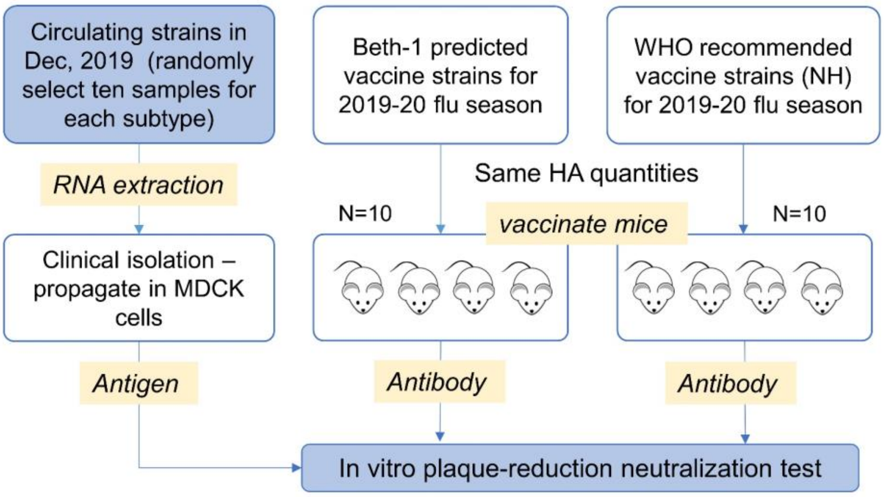


**Extended Data Fig. 5 | Experimental design of in vivo experiment.** Batches of female BALB/c mice aged at 6-8 weeks were immunized intraperitoneally by recombinant influenza virus with the predicted HA and NA genes by the current-system and beth-1 were. The levels of neutralizing antibodies in antisera were determined by PRNT using clinical isolates of the pH1N1 or H3 viruses.

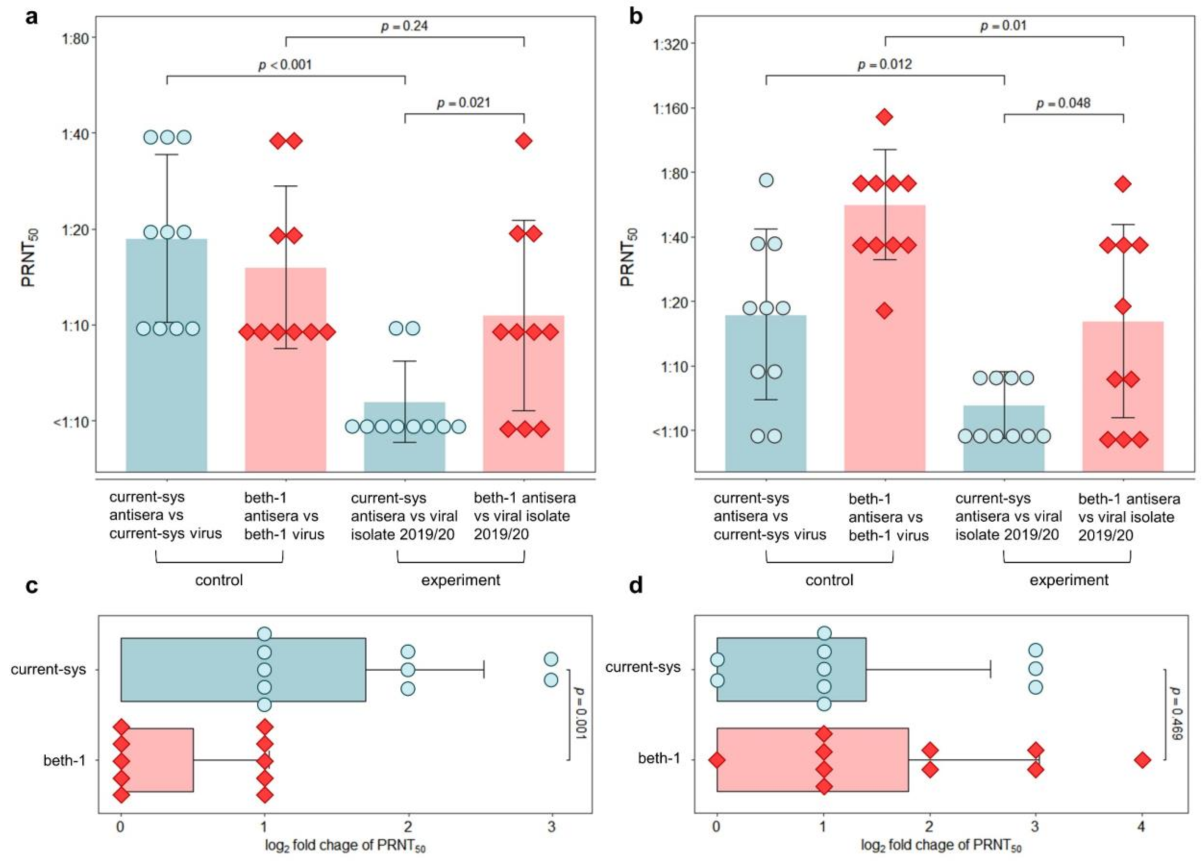


**Extended Data Fig. 6 | Neutralization of antibodies induced by the predicted strains for the 2019/20 season in Hong Kong. a**, neutralization for pH1N1 virus. **b**, neutralization for H3N2 virus. **c**, pH1N1, titer fold-change. **d**, H3N2, titer fold-change. The neutralizing effect of antibodies elicited by predicted strains in mouse model was evaluated by $PRNT_{50}$ titer against virus isolates and compared to those elicited by strains predicted by the current system. Control groups: antibodies elicited by predicted strain neutralizing the predicted strain. Experimental group: antibodies elicited by predicted strains neutralizing viral isolates in Hong Kong 2019/20. Fold-change: the ratio of titer of control group to titer of experimental group. Beth-1 predicted strains (red color) generated neutralization significantly higher (*t*-test *p*-value = 0.001) for the pH1N1 and non-inferior for the H3N2.

**Extended Data Table 1 | Effective mutations and predictor codon sites.** Effective mutation sites (EMs) emerged in each season are listed. **Boldface**: epitope sites. Underline: predictor codon sites. The codon sites of pH1N1 are in H3 numbering.

| Flu season | HA protein | NA protein |
|---|---|---|
| *Subtype: pH1N1* | | |
| 2011/12 | **188**, 453 | 241, 369 |
| 2012/13 | 2, 92, 146, 200, 285, 501 | 44, 200, 232 |
| 2013/14 | 237 | - |
| 2014/15 | **166**, 259, 271 | 34, 40, 321, 432 |
| 2015/16 | 7 | 13, **264**, **270**, 314 |
| 2016/17 | **165**, 219 | - |
| 2017/18 | **82**, 218, 297 | 188, **449** |
| 2018/19 | **167** | 77, 81 |
| 2019/20 | 186 | 51, 74, 416, 452 |
| 2020/21 | 133, 134, **159**, **164**, **190**, **192**, 253, 262, 508 | 19, 52, 66, 222, **389** |
| 2021/22 | 169, 205, 368 | 432 |
| 2022/23 | 101 | 469 |
| *Subtype: H3N2* | | |
| 2002/03 | 25, 75, 83, **155**, **156**, 183, **186**, 202, 222, **309**, 500, 530 | 30, 42, 143, 216, 332, 385 |
| 2003/04 | 479 | - |
| 2004/05 | **126**, **138**, **145**, **159**, **189**, **227** | 18, 40, 93, 172, **199**, 265, 399, 432, 437 |
| 2005/06 | **193**, **226**, 326, 361, 484 | **221** |
| 2006/07 | 6, **157**, 375, 450 | 4, 5, 43, 194, 310, 315, **370** |
| 2007/08 | **142** | 86, **150**, 151, 296, 335 |
| 2009/10 | - | - |
| 2010/11 | **62**, **158**, **213** | - |
| 2011/12 | **53**, **94**, **212**, **230**, **280** | 210, 367, 369, 464 |
| 2012/13 | **45**, **48**, **198**, 223, **278**, **312** | 81, 402 |
| 2013/14 | - | - |
| 2014/15 | **128** | 392 |
| 2015/16 | 3, **160**, **311** | - - |
| 2016/17 | **171**, 406, 489 | 245, 247, 339, 380, 468 |
| 2017/18 | **261** | - |
| 2018/19 | **121**, **135** | **220**, 303, **329** |
| 2019/20 | **137**, 529 | 315, **344** |
| 2020/21 | - | 469 |
| 2021/22 | **83**, **131**, **164**, **186**, **190**, 195, 522 | 62, 346, 463, 465 |
| 2022/23 | 104, **276** | - |

**Extended Data Table 2 | Sequence sample size.** Complete genetic sequences of influenza virus pH1N1 and H3N2 were retrieved from the Global Initiative on Sharing all Influenza Data (GISAID). The total number of sequences used in analysis is 50,285 for hemagglutinin (HA) and 45,297 for Neuraminidase (NA).

| Flu season | **pH1N1** | | **H3N2** | |
|---|---|---|---|---|
| | HA | NA | HA | NA |
| 1999/00 | - | - | 87 | 87 |
| 2000/01 | - | - | 25 | 23 |
| 2001/02 | - | - | 112 | 110 |
| 2002/03 | - | - | 52 | 50 |
| 2003/04 | - | - | 236 | 237 |
| 2004/05 | - | - | 164 | 161 |
| 2005/06 | - | - | 73 | 73 |
| 2006/07 | - | - | 86 | 84 |
| 2007/08 | - | - | 88 | 85 |
| 2008/09 | - | - | 434 | 426 |
| 2009/10 | 1817 | 922 | 129 | 128 |
| 2010/11 | 226 | 167 | 325 | 295 |
| 2011/12 | 51 | 47 | 546 | 454 |
| 2012/13 | 184 | 163 | 454 | 305 |
| 2013/14 | 461 | 448 | 626 | 508 |
| 2014/15 | 289 | 288 | 1733 | 1329 |
| 2015/16 | 1335 | 1310 | 852 | 789 |
| 2016/17 | 551 | 551 | 2907 | 2477 |
| 2017/18 | 1220 | 1122 | 2179 | 1670 |
| 2018/19 | 2063 | 1714 | 2548 | 2190 |
| 2019/20 | 1566 | 1193 | 1875 | 1671 |
| 2020/21[a] | 122 | 111 | 494 | 469 |
| 2021/22[a] | 1358 | 1360 | 15476 | 15074 |
| 2022/23 | 1949 | 1864 | 5592 | 5342 |
| **Sum** | **13,192** | **11,260** | **37,093** | **34,037** |

[a] In these periods, all available sequences in the Northern Hemisphere (NH) were retrieved. Other than these epidemics, all available sequences from ten geographical regions in NH were retrieved, include North America (New York State, California State, Canada), Europe (United Kingdom, Germany, France), and Asia (Hong Kong SAR, South China provinces, Japan, Singapore).

### Extended Data Table 3 | Summary table of predicted strains

| Flu season | WHO recommended vaccine strains[a] | beth-1 strain (two-protein model) (sequence collection time) | Answer strain[c] |
|---|---|---|---|
| *Subtype: pH1N1* | | | |
| 2012/13 | A/California/07/2009 | A/India/Nsk12388/2012 (2012-01-19) | A/India/2080/2012 – like virus |
| 2013/14 | A/California/07/2009 | A/Singapore/GP325/2013 (2013-02-07) | A/Hong Kong/6015/2013 – like virus |
| 2014/15 | A/California/07/2009 | A/Singapore/KK107/2014 (2014-01-06) | A/Norway/2432/2014 – like virus |
| 2015/16 | A/California/07/2009 | A/India/P153017/2015 (2015-02-23) | A/Hong Kong/15587/2015 – like virus |
| 2016/17 | A/California/07/2009 | A/Prachuapkhirikhan/42/2016 (2016-02-29) | A/Singapore/SGH0769/2016 – like virus |
| 2017/18 | A/Michigan/45/2015 | A/Phra Nakhon Si Ayutthaya/396/2016 (2016-11-01) | A/Oman/5640/2017 – like virus |
| 2018/19 | A/Michigan/45/2015 | A/Hong Kong/4989/2017 (2017-12-17) | A/Lipetsk/1V/2018 – like virus |
| 2019/20[b] | A/Brisbane/02/2018 | A/Hong Kong/98/2019 (2019-01-01)<br>A/Hong Kong/2655/2019 (2019-06-18) | A/Bahrain/578/2019 – like virus |
| 2020/21 | A/Guangdong-Maonan/SWL1536/2019 | A/Hong Kong/4380/2019 (2019-11-27) | A/India/PUN-NIV323483/2021 – like virus |
| 2021/22 | A/Victoria/2570/2019 | A/Nagasaki/8/2020 (2020-10-27) | A/Netherlands/10201/2021 – like virus |
| 2022/23 | A/Victoria/2570/2019 | A/Bangladesh/3210910005/2021 (2021-09-07) | A/Denmark/3091/2022 – like virus |
| *Subtype: H3N2* | | | |
| 2002/03 | A/Moscow/10/99 | A/Hong Kong/CUHK5250/2002 (2002-01-10) | A/Denmark/32/03 – like virus |
| 2003/04 | A/Moscow/10/99 | A/Hong Kong/CUHK5316/2003 (2003-01-10) | A/New York/54/2003 – like virus |
| 2004/05 | A/Fujian/411/2002 | A/Singapore/NHRC0012/2003 (2003-10-13) | A/Hong Kong/CUHK10026/2005 – like virus |
| 2005/06 | A/California/07/2004 | A/Guangdong/03/2005 (2005-01-01) | A/Illinois/NHRC0002/2005 – like virus |
| 2006/07 | A/Wisconsin/67/2005 | A/Thailand/CU259/2006 (2006-01-01) | A/Singapore/68Q/2007 – like virus |
| 2007/08 | A/Wisconsin/67/2005 | A/Beijing/Xicheng1272/2006 (2006-12-11) | A/Florida/UR07-0101/2008 – like virus |
| 2008/09 | A/Brisbane/10/2007 | A/Thailand/CU379/2008 (2008-01-01) | A/California/VRDL375/2009 – like virus |
| 2009/10 | A/Brisbane/10/2007 | A/Thailand/CU379/2008 (2008-01-01) | A/Florida/26/2009 – like virus |
| 2010/11 | A/Perth/16/2009 | A/Hong Kong/H090-783-V10/2009 (2009-08-07) | A/Kentucky/12/2010 – like virus |
| 2011/12 | A/Perth/16/2009 | A/Singapore/C2011.027V/2011 (2011-01-10) | A/Montana/08/2011 – like virus |
| 2012/13 | A/Victoria/361/2011 | A/Hunan-Kifu/1166/2012 (2012-01-01) | A/Arizona/09/2012 – like virus |
| 2013/14 | A/Victoria/361/2011 | A/Hong Kong/8147/2012 (2012-06-26) | A/South Carolina/03/2014 – like virus |
| 2014/15 | A/Texas/50/2012 | A/Singapore/H2013.403/2013 (2013-05-27) | A/Texas/38/2014 – like virus |
| 2015/16 | A/Switzerland/9715293/2013 | A/Shandong-Linzi/323/2014 (2014-12-01) | A/Colorado/15/2015 – like virus |
| 2016/17 | A/Hong Kong/4801/2014 | A/Chiang Mai/54/2016 (2016-02-13) | A/Singapore/TT0348/2017 – like virus |
| 2017/18 | A/Hong Kong/4801/2014 | A/Singapore/KK0164/2017 (2017-02-20) | A/Pennsylvania/263/2017 – like virus |
| 2018/19 | A/Singapore/INFIMH-16-0019/2016 | A/Wuhan/2785/2018 (2018-02-01) | A/Kyiv/537/2018 – like virus |
| 2019/20 | A/Kansas/14/2017 | A/Hong Kong/2926/2018 (2018-11-26)<br>A/Hong Kong/2678/2019 (2019-06-18) | A/Hungary/9/2020 – like virus |
| 2020/21 | A/Hong Kong/2671/2019 | A/Hong Kong/4435/2019 (2019-11-29) | A/Michigan/03/2021 – like virus |
| 2021/22 | A/Cambodia/e0826360/2020 | A/Bangladesh/0004/2020 (2020-09-06) | A/Michigan/08/2021 – like virus |
| 2022/23 | A/Darwin/6/2021 | A/Arsal/5773/2021 (2021-09-13) | A/England/224040474/2022 – like virus |

[a] The WHO recommended vaccine for Northern Hemisphere were collected from influenza vaccine sheet of WHO, accessed via https://www.who.int/teams/global-influenza-programme/vaccines/who-recommendations, and the sequences were retrieved from GISAID [25].

[b] Shaded rows: Seasons in prospective study.

[c] The answer strain is the closest wild-type strain to the consensus strain obtained from the observed circulating virus in the target season.

**Extended Data Table 4 | Summary table of predicted strains by the LBI and beth-1 (single protein)**

| Flu season | beth-1 vaccine strain (HA) | beth-1 vaccine strain (NA) | LBI (HA)[a] | LBI (NA) |
|---|---|---|---|---|
| *Subtype: pH1N1* | | | | |
| 2012/13 | A/India/Nsk12388/2012 | A/Srinigar/827/2011 | A/Florida/19/2011 | A/Florida/27/2011 |
| 2013/14 | A/Singapore/GP325/2013 | A/Singapore/GP325/2013 | A/Taiwan/989/2012 | A/Lithuania/5509/2013 |
| 2014/15 | A/Singapore/KK107/2014 | A/Singapore/KK1064/2013 | A/Haiti/2036/2013 | A/YOKOHAMA/162/2013 |
| 2015/16 | A/India/3388/2015 | A/India/P153017/2015 | A/Berlin/87/2014 | A/Trencin/298/2015 |
| 2016/17 | A/Prachuapkhirikhan/42/2016 | A/Prachuapkhirikhan/42/2016 | A/Kyrgyzstan/71/2015 | A/Wisconsin/21/2016 |
| 2017/18 | A/Phra_Nakhon_Si_Ayutthaya/396/2016 | A/Singapore/TT0169/2017 | A/Ohio/07/2017 | A/Shanghai-Minxing/SWL113/2017 |
| 2018/19 | A/Singapore/GP0014/2018 | A/Singapore/EN0138/2018 | A/Washington/299/2017 | A/Jeju/836/2017 |
| *Subtype: H3N2* | | | | |
| 2002/03 | A/Hong Kong/CUHK5250/2002 | A/Hong Kong/CUHK5250/2002 | A/New York/92/2002 | A/Texas/NHRC0001/2002 |
| 2003/04 | A/Hong Kong/CUHK5316/2003 | A/Hong Kong/CUHK6383/2003 | A/Hong Kong/CUHK33047/2002 | A/Genoa/1/2002 |
| 2004/05 | A/Hong Kong/CUHK74438/2003 | A/Hong Kong/HKU1/2004 | A/Thailand/Siriraj-02/2003 | A/New Jersey/NHRC0001/2003 |
| 2005/06 | A/Guangdong/04/2005 | A/Hong Kong/CUHK7047/2005 | A/New York/469/2004 | A/South Carolina/NHRC0001/2005 |
| 2006/07 | A/Thailand/CU259/2006 | A/Thailand/CU259/2006 | A/New York/1001/2005 | A/New York/1012/2006 |
| 2007/08 | A/Beijing/Xicheng1272/2006 | A/Hong Kong/CUHK53005/2006 | A/Vietnam/19/2007 | A/Honduras/45/2006 |
| 2008/09 | A/Thailand/CU379/2008 | A/Thailand/CU379/2008 | A/Singapore/65/2007 | A/Louisiana/05/2008 |
| 2009/10 | A/Hong Kong/40/2009 | A/Singapore/619/2008 | A/Korea/WRAIR1391P/2009 | A/North Dakota/02/2008 |
| 2010/11 | A/Hong Kong/H090-689-V2 | A/SURAT_THANI/116/2010 | A/Oregon/08/2009 | A/Pennsylvania/02/2010 |
| 2011/12 | A/Singapore/KK114/2011 | A/Singapore/H2011.140C/2011 | A/Pennsylvania/10/2010 | A/Ontario/P21548/2010 |
| 2012/13 | A/Hunan-Kifu/1166/2012 | A/China/89/2012 | A/Singapore/KK219/2011 | A/Paris/2100/2011 |
| 2013/14 | A/Delhi/764/2013 | A/Hangzhou/A155/2013 | A/Helsinki/942/2013 | A/Houston/JMM 158/2013 |
| 2014/15 | A/Anhui-Yaohai/1704/2013 | A/Singapore/GP415/2014 | A/Florida/21/2013 | A/Norway/3303/2013 |
| 2015/16 | A/Shaanxi-Baota/1661/2014 | A/China/60376/2014 | A/OSAKA/63/2014 | A/Tennessee/16/2014 |
| 2016/17 | A/Shanxi-Yaodou/1327/2016 | A/Singapore/INFTT-16-0224/2016 | A/Ontario/RV2425/2015 | A/Mexico/92/2016 |
| 2017/18 | A/Singapore/GP0269/2017 | A/Singapore/TT0096/2017 | A/Singapore/MOH0107/2016 | A/Champagne Ardenne/628/2017 |
| 2018/19 | A/Wuhan/2785/2018 | A/Singapore/TT0256/2018 | A/Zhejiang-Yongkang/1978/2017 | A/Ireland/61097/2017 |

[a] When the highest LBI score corresponds to multiple strains in a season, one strain is randomly picked.

**Extended Data Table 5 | Summary of prediction performance in retroactive validation by alternative methods for the HA and NA genes of pH1N1 and H3N2**

| Subtype | Prediction seasons | Gene | Average number of amino acids mismatch[a,b] (SD) | | | | | | |
|---|---|---|---|---|---|---|---|---|---|
| | | | current-system | beth-1 (HA+NA) | answer strain | LBI (HA) | beth-1 (HA) | LBI (NA) | beth-1 (NA) |
| pH1N1 | 2012/13 - 2018/19 | HA full-length | 12.8 (4.4) | 5.1 (2.8) | 3.0 (1.2) | 5.7 (2.5) | 5.1 (2.8) | - | - |
| | | NA full-length | 11.6 (4.4) | 4.4 (1.6) | 2.9 (0.9) | - | - | 6.4 (2.1) | 3.9 (1.5) |
| | | HA epitope | 3.4 (0.8) | 1.2 (0.6) | 0.7 (0.4) | 1.6 (0.9) | 1.2 (0.6) | - | - |
| | | NA epitope | 1.2 (0.8) | 0.5 (0.4) | 0.4 (0.2) | - | - | 1.2 (1.1) | 0.5 (0.4) |
| H3N2 | 2002/03 - 2018/19 | HA full-length | 11.7 (5.1) | 7.5 (2.2) | 5.4 (1.7) | 9.5 (4.7) | 7.5 (2.2) | - | - |
| | | NA full-length | 7.7 (2.5) | 5.7 (3.0) | 4.3 (2.2) | - | - | 7.3 (3.6) | 5.3 (2.3) |
| | | HA epitope | 7.7 (3.4) | 5.1 (1.7) | 3.5 (1.1) | 6.1 (2.9) | 5.1 (1.7) | - | - |
| | | NA epitope | 1.0 (0.8) | 0.6 (0.5) | 0.4 (0.4) | - | - | 0.7 (0.6) | 0.5 (0.5) |

[a] Average amino acids mismatch calculated from pair-wise mismatch between circulating strains against vaccine strain. Epidemic seasons predicted: for pH1N1, 7 seasons from 2012/13 to 2018/19; and for H3N2, 17 seasons from 2002/03 to 2018/19. Season-to-season breakdown is in **Extended Data Table 6-9**.

[b] Ten geographical regions evaluated in Northern Hemisphere, covering North America (New York State, California State, Canada), Europe (United Kingdom, Germany, France), and Asia (Hong Kong SAR, South China provinces, Japan, Singapore). Genetic mismatch breakdown by geographical region is in **Extended Data Table 10-13**.

**Extended Data Table 6 | Prediction performance in retrospective data by epidemic season for HA of pH1N1**

| Epidemic season (sample size) | Average number of mismatch on HA sequence (out of 566 codons) (amino acids, AAs) | | | | Average number of mismatch[a] on HA epitope (out of 50 sites) (AAs) | | | |
|---|---|---|---|---|---|---|---|---|
| | current-system (SD) | beth-1 (HA+NA) (SD) | LBI (HA) (SD) | beth-1 (HA) (SD) | current-system (SD) | beth-1 (HA+NA) (SD) | LBI (HA) (SD) | beth-1 (HA) (SD) |
| 2012/13 (N=184) | 13.0 (0.6) | 10.4 (1.1) | 10.2 (1.1) | 10.4 (1.1) | 3.0 (0.3) | 1.0 (0.3) | 2.8 (0.3) | 1.0 (0.3) |
| 2013/14 (N=461) | 12.9 (0.6) | 4.3 (0.4) | 5.4 (0.4) | 4.3 (0.4) | 3.2 (0.2) | 1.2 (0.2) | 1.2 (0.2) | 1.2 (0.2) |
| 2014/15 (N=289) | 14.1 (1.1) | 2.2 (1.1) | 2.2 (1.1) | 2.2 (1.1) | 3.4 (0.4) | 0.4 (0.4) | 0.4 (0.4) | 0.4 (0.4) |
| 2015/16 (N=1,335) | 16.8 (0.5) | 3.3 (1.1) | 4.9 (0.6) | 3.3 (1.1) | 4.0 (0.2) | 1.0 (0.2) | 1.0 (0.2) | 1.0 (0.2) |
| 2016/17 (N=551) | 18.6 (0.8) | 2.9 (0.6) | 3.9 (0.6) | 2.9 (0.6) | 5.0 (0.6) | 1.1 (0.5) | 1.1 (0.5) | 1.1 (0.5) |
| 2017/18 (N=1,220) | 5.9 (0.9) | 6.9 (0.9) | 6.9 (0.9) | 6.9 (0.9) | 2.4 (0.2) | 2.4 (0.2) | 2.4 (0.2) | 2.4 (0.2) |
| 2018/19 (N=2,063) | 8.4 (0.9) | 5.4 (0.9) | 6.4 (0.9) | 5.4 (0.9) | 3.2 (0.3) | 1.2 (0.3) | 2.2 (0.3) | 1.2 (0.3) |

[a] Average amino acids mismatch against all available circulating strains in respective epidemic season in ten geographical regions.

**Extended Data Table 7 | Prediction performance in retrospective data by epidemic season for NA of pH1N1**

| Epidemic season (sample size) | Average number of mismatch on HA sequence (out of 469 codons) (AAs) | | | | Average number of mismatch[a] on HA epitope (out of 30 sites) (AAs) | | | |
|---|---|---|---|---|---|---|---|---|
| | current-system (SD) | beth-1 (HA+NA) (SD) | LBI (NA) (SD) | beth-1 (NA) (SD) | current-system (SD) | beth-1 (HA+NA) (SD) | LBI (NA) (SD) | beth-1 (NA) (SD) |
| 2012/13 (N=163) | 7.9 (0.7) | 4.5 (1.1) | 4.6 (0.7) | 2.5 (1.0) | 0.2 (0.1) | 0.2 (0.1) | 0.2 (0.1) | 0.2 (0.1) |
| 2013/14 (N=448) | 11.6 (0.6) | 5.8 (0.5) | 6.7 (0.5) | 5.8 (0.5) | 0.3 (0.3) | 0.3 (0.3) | 0.3 (0.3) | 0.3 (0.3) |
| 2014/15 (N=288) | 14.2 (0.8) | 5.7 (1.1) | 5.6 (0.8) | 4.7 (1.1) | 0.9 (0.4) | 0.9 (0.4) | 0.9 (0.4) | 0.9 (0.4) |
| 2015/16 (N=1,310) | 15.5 (0.5) | 1.7 (1.2) | 7.8 (0.5) | 1.7 (1.2) | 2.0 (0.1) | 0.2 (0.1) | 2.0 (0.1) | 0.2 (0.1) |
| 2016/17 (N=551) | 17.6 (0.8) | 2.9 (0.8) | 3.8 (0.7) | 2.9 (0.8) | 2.5 (0.4) | 0.5 (0.4) | 1.5 (0.3) | 0.5 (0.4) |
| 2017/18 (N=1,122) | 6.0 (0.5) | 6.0 (0.5) | 9.9 (0.5) | 5.0 (0.5) | 1.2 (0.2) | 1.2 (0.2) | 3.2 (0.2) | 1.2 (0.2) |
| 2018/19 (N=1,714) | 8.1 (0.8) | 4.5 (0.6) | 6.4 (0.7) | 4.5 (0.6) | 1.4 (0.2) | 0.4 (0.2) | 0.4 (0.2) | 0.4 (0.2) |

[a] Average amino acids mismatch against all available circulating strains in respective epidemic season in ten geographical regions.

**Extended Data Table 8 | Prediction performance in retrospective data by epidemic season for HA of H3N2**

| Epidemic season (sample size) | Average mismatch on HA sequence (out of 566 codons) (AAs) | | | | Average mismatch[a] on HA epitope (out of 131 sites) (AAs) | | | |
|---|---|---|---|---|---|---|---|---|
| | current-system (SD) | beth-1 (HA+NA) (SD) | LBI (HA) (SD) | beth-1 (HA) (SD) | current-system (SD) | beth-1 (HA+NA) (SD) | LBI (HA) (SD) | beth-1 (HA) (SD) |
| 2002/03 (N=52) | 23.3 (2.9) | 9.0 (1.5) | 18.1 (4.1) | 9.0 (1.5) | 14.6 (1.9) | 5.0 (0.3) | 8.9 (2.0) | 5.0 (0.3) |
| 2003/04 (N=236) | 25.1 (0.8) | 4.7 (1.4) | 4.8 (1.3) | 4.7 (1.4) | 16.5 (0.8) | 3.2 (1.0) | 4.2 (1.0) | 3.2 (1.0) |
| 2004/05 (N=164) | 11.8 (3.5) | 6.7 (3.8) | 6.8 (3.8) | 6.7 (3.8) | 7.9 (2.4) | 4.5 (2.5) | 4.6 (2.4) | 4.5 (2.5) |
| 2005/06 (N=73) | 10.3 (4.7) | 6.7 (6.1) | 7.5 (4.7) | 6.7 (6.1) | 6.3 (2.7) | 3.1 (3.0) | 3.5 (2.6) | 3.1 (3.0) |
| 2006/07 (N=86) | 10.7 (2.7) | 7.4 (2.4) | 6.9 (2.9) | 7.4 (2.4) | 6.3 (1.4) | 4.6 (1.3) | 3.4 (1.5) | 4.6 (1.3) |
| 2007/08 (N=88) | 11.0 (0.7) | 9.1 (0.8) | 7.9 (0.7) | 9.1 (0.8) | 6.6 (0.5) | 5.6 (0.5) | 5.4 (0.6) | 5.6 (0.5) |
| 2008/09 (N=434) | 8.3 (1.7) | 4.3 (1.7) | 6.3 (1.7) | 4.3 (1.7) | 4.6 (1.5) | 3.6 (1.6) | 4.6 (1.5) | 3.6 (1.6) |
| 2009/10 (N=129) | 11.7 (0.6) | 7.7 (0.6) | 17.6 (0.6) | 7.7 (0.6) | 7.6 (0.6) | 6.6 (0.6) | 10.6 (0.6) | 6.6 (0.6) |
| 2010/11 (N=325) | 9.5 (0.7) | 9.4 (0.8) | 9.0 (0.7) | 9.4 (0.8) | 7.9 (0.6) | 7.9 (0.7) | 7.4 (0.6) | 7.9 (0.7) |
| 2011/12 (N=546) | 11.0 (0.7) | 10.1 (1.8) | 7.2 (0.8) | 10.1 (1.8) | 8.6 (0.6) | 7.7 (1.6) | 4.8 (0.7) | 7.7 (1.6) |
| 2012/13 (N=454) | 6.9 (0.8) | 4.6 (1.1) | 5.9 (0.8) | 4.6 (1.1) | 3.8 (0.7) | 2.3 (0.8) | 3.8 (0.7) | 2.3 (0.8) |
| 2013/14 (N=626) | 9.3 (0.9) | 5.5 (0.8) | 18.8 (1) | 5.5 (0.8) | 5.2 (0.5) | 3.3 (0.5) | 13.8 (0.3) | 3.3 (0.5) |
| 2014/15 (N=1,733) | 13.1 (0.5) | 9.5 (0.8) | 10.3 (0.7) | 9.5 (0.8) | 9.6 (0.3) | 6.0 (0.5) | 6.0 (0.5) | 6.0 (0.5) |
| 2015/16 (N=852) | 11.5 (2.6) | 10.9 (2.2) | 5.8 (1.2) | 10.9 (2.2) | 7.3 (1.6) | 7.3 (1.6) | 3.8 (1.1) | 7.3 (1.6) |
| 2016/17 (N=2,907) | 6.3 (0.8) | 5.8 (1.1) | 5.8 (1.1) | 5.8 (1.1) | 4.3 (0.8) | 4.1 (0.9) | 4.1 (0.9) | 4.1 (0.9) |
| 2017/18 (N=2,179) | 6.8 (1.4) | 6.0 (2.8) | 7.6 (0.4) | 6.0 (2.8) | 5.4 (0.9) | 4.7 (2.4) | 5.6 (0.5) | 4.7 (2.4) |
| 2018/19 (N=2,548) | 12.0 (2.6) | 10.4 (3.3) | 15.1 (2.6) | 10.4 (3.3) | 7.4 (1.2) | 6.7 (1.9) | 8.7 (2.4) | 6.7 (1.9) |

[a] Average amino acids mismatch against all available circulating strains in respective epidemic season, averaging the mismatch in ten geographical regions.

**Extended Data Table 9 | Prediction performance in retrospective data by epidemic season for NA of H3N2**

| Epidemic season (sample size) | Average number of mismatch on NA sequence (out of 469 codons) (AAs) | | | | Average number of mismatch[a] on NA epitope (out of 12 sites) (AAs) | | | |
|---|---|---|---|---|---|---|---|---|
| | current-system (SD) | beth-1 (HA+NA) (SD) | LBI (NA) (SD) | beth-1 (NA) (SD) | current-system (SD) | beth-1 (HA+NA) (SD) | LBI (NA) (SD) | beth-1 (NA) (SD) |
| 2002/03 (N=52) | 10.5 (1.1) | 4.5 (4.3) | 12.3 (3.3) | 4.5 (4.3) | 0.3 (0.3) | 0.3 (0.3) | 0.3 (0.3) | 0.3 (0.3) |
| 2003/04 (N=236) | 11.3 (1.0) | 8.4 (5.6) | 10.9 (4.9) | 8.4 (5.6) | 1.0 (0.2) | 1.0 (0.2) | 1.0 (0.2) | 1.0 (0.2) |
| 2004/05 (N=164) | 6.4 (2.5) | 4.1 (2.1) | 16.0 (1.6) | 4.1 (2.1) | 1.8 (0.4) | 0.8 (0.4) | 0.8 (0.4) | 0.8 (0.4) |
| 2005/06 (N=73) | 5.3 (4.9) | 4.9 (5.0) | 5.3 (4.9) | 5.3 (4.9) | 0.8 (1.3) | 0.8 (1.3) | 0.8 (1.3) | 0.8 (1.3) |
| 2006/07 (N=86) | 8.3 (1.5) | 9.0 (3.2) | 7.3 (1.5) | 9.0 (3.2) | 1.5 (0.4) | 1.0 (0.5) | 1.5 (0.4) | 1.0 (0.5) |
| 2007/08 (N=88) | 10.6 (0.7) | 14.8 (0.8) | 11.9 (0.9) | 10.5 (0.7) | 2.1 (0.1) | 1.1 (0.1) | 2.1 (0.1) | 0.1 (0.1) |
| 2008/09 (N=434) | 4.7 (0.7) | 1.8 (0.8) | 5.6 (0.6) | 1.8 (0.8) | 0.1 (0.1) | 0.1 (0.1) | 0.1 (0.1) | 0.1 (0.1) |
| 2009/10 (N=129) | 7.4 (1.6) | 4.4 (1.6) | 7.0 (2.2) | 4.4 (1.6) | 0.0 (0.1) | 0.0 (0.1) | 0.0 (0.1) | 0.0 (0.1) |
| 2010/11 (N=325) | 5.4 (1.2) | 5.4 (1.2) | 3.6 (1.1) | 5.4 (1.2) | 0.1 (0.1) | 0.1 (0.1) | 0.1 (0.1) | 0.1 (0.1) |
| 2011/12 (N=546) | 6.9 (0.4) | 4.9 (0.5) | 3.9 (0.5) | 4.9 (0.5) | 0.1 (0.1) | 0.1 (0.1) | 0.1 (0.1) | 0.1 (0.1) |
| 2012/13 (N=454) | 3.9 (0.7) | 2.9 (0.7) | 7.4 (0.7) | 1.9 (0.7) | 1.1 (0.2) | 0.2 (0.2) | 0.9 (0.1) | 0.2 (0.2) |
| 2013/14 (N=626) | 5.8 (0.5) | 3.9 (0.5) | 3.9 (0.5) | 3.9 (0.5) | 1.7 (0.3) | 0.7 (0.3) | 0.7 (0.3) | 0.7 (0.3) |
| 2014/15 (N=1,733) | 6.6 (0.5) | 5.1 (0.2) | 4.1 (0.3) | 4.8 (0.4) | 2.0 (0.1) | 0.2 (0.1) | 0.2 (0.1) | 0.2 (0.1) |
| 2015/16 (N=852) | 7.1 (0.8) | 7.1 (0.8) | 4.1 (1.4) | 6.5 (0.5) | 0.2 (0.1) | 0.2 (0.1) | 0.2 (0.1) | 0.2 (0.1) |
| 2016/17 (N=2,907) | 10.1 (0.8) | 3.8 (1.4) | 7.4 (0.8) | 3.8 (1.4) | 0.6 (0.3) | 0.6 (0.3) | 0.6 (0.3) | 0.6 (0.3) |
| 2017/18 (N=2,179) | 12.1 (0.3) | 4.7 (0.4) | 4.7 (0.6) | 4.4 (0.4) | 1.3 (0.2) | 1.3 (0.2) | 0.6 (0.3) | 1.3 (0.2) |
| 2018/19 (N=2,548) | 8.3 (0.9) | 8.2 (2.4) | 7.8 (0.7) | 6.2 (2.3) | 2.4 (0.2) | 1.5 (0.7) | 1.7 (0.2) | 1.5 (0.7) |

[a] Average amino acids mismatch against all available circulating strains in respective epidemic season in ten geographical regions.

**Extended Data Table 10 | Prediction performance in retrospective data by geographical regions for HA of pH1N1**

| Country/region[a] (sample size) | Average mismatch on HA sequence (out of 566 codons) (AAs) | | | | Average mismatch[b] on HA epitope (out of 50 sites) (AAs) | | | |
|---|---|---|---|---|---|---|---|---|
| | current-system (SD) | beth-1 (HA+NA) (SD) | LBI (HA) (SD) | beth-1 (HA) (SD) | current-system (SD) | beth-1 (HA+NA) (SD) | LBI (HA) (SD) | beth-1 (HA) (SD) |
| NY (N=134) | 13.6 (4.3) | 5.6 (2.7) | 6.4 (2.6) | 5.6 (2.7) | 3.7 (1.0) | 1.5 (0.4) | 1.8 (0.7) | 1.5 (0.4) |
| CA (N=353) | 12.4 (4.4) | 4.8 (3.1) | 5.3 (2.6) | 4.8 (3.1) | 3.3 (0.9) | 1.1 (0.6) | 1.4 (0.8) | 1.1 (0.6) |
| CAN (N=373) | 12.8 (4.8) | 5.4 (3.1) | 5.7 (1.9) | 5.4 (3.1) | 3.5 (1.3) | 1.3 (0.8) | 1.7 (0.8) | 1.3 (0.8) |
| UK (N=1,197) | 12.5 (4.8) | 4.8 (3.0) | 5.2 (2.1) | 4.8 (3.0) | 3.5 (1.0) | 1.2 (0.6) | 1.5 (0.7) | 1.2 (0.6) |
| GER (N=173) | 12.2 (4.5) | 4.4 (3.0) | 5.0 (2.6) | 4.4 (3.0) | 3.3 (0.7) | 1.0 (0.6) | 1.4 (1.0) | 1.0 (0.6) |
| FR (N=767) | 12.4 (4.5) | 4.5 (3.0) | 5.3 (3.1) | 4.5 (3.0) | 3.3 (0.9) | 1.1 (0.7) | 1.5 (0.9) | 1.1 (0.7) |
| HK (N=334) | 12.7 (4.5) | 4.9 (3.1) | 5.5 (2.6) | 4.9 (3.1) | 3.4 (0.7) | 1.1 (0.7) | 1.5 (1.1) | 1.1 (0.7) |
| SCP (N=823) | 12.5 (4.3) | 5.5 (3.2) | 5.8 (2.8) | 5.5 (3.2) | 3.2 (0.6) | 1.0 (0.7) | 1.4 (1.1) | 1.0 (0.7) |
| JP (N=1,292) | 14 (4.2) | 6.1 (2.9) | 6.9 (2.9) | 6.1 (2.9) | 3.7 (0.7) | 1.4 (0.8) | 1.8 (1.1) | 1.4 (0.8) |
| SG (N=657) | 12.9 (4.5) | 4.7 (2.2) | 5.8 (2.8) | 4.7 (2.2) | 3.4 (0.8) | 1.2 (0.7) | 1.6 (1.0) | 1.2 (0.7) |
| **Average (SD)** | **12.8 (4.4)** | **5.1 (2.8)** | **5.7 (2.5)** | **5.1 (2.8)** | **3.4 (0.8)** | **1.2 (0.6)** | **1.6 (0.9)** | **1.2 (0.6)** |

[a] Abbreviations: NY: New York State, CA: California, CAN: Canada, UK: United Kingdom, FR: France, HK: Hong Kong SAR, SCP: South China Provinces, JP: Japan, SG: Singapore

[b] Average amino acids mismatch against all available circulating strains in epidemic seasons from 2012/13 to 2018/19 in respective geographical regions.

**Extended Data Table 11 | Prediction performance in retrospective data by geographical regions for NA of pH1N1**

| Country/region[a] (sample size) | Average number of mismatch on NA sequence (out of 469 codons) (AAs) | | | | Average number of mismatch[b] on NA epitope (out of 30 sites) (AAs) | | | |
|---|---|---|---|---|---|---|---|---|
| | current-system (SD) | beth-1 (HA+NA) (SD) | LBI (NA) (SD) | beth-1 (NA) (SD) | current-system (SD) | beth-1 (HA+NA) (SD) | LBI (NA) (SD) | beth-1 (NA) (SD) |
| NY (N=134) | 11.4 (4.6) | 3.9 (1.7) | 6.1 (1.9) | 3.4 (1.6) | 1.1 (0.8) | 0.5 (0.6) | 1.1 (1.2) | 0.5 (0.6) |
| CA (N=353) | 11.1 (4.6) | 3.8 (1.8) | 6 (2.0) | 3.2 (1.8) | 1.1 (0.9) | 0.4 (0.5) | 1.1 (1.2) | 0.4 (0.5) |
| CAN (N=373) | 11.7 (5.1) | 4.7 (2.2) | 6.4 (2.0) | 4.2 (2.0) | 1.4 (0.9) | 0.7 (0.4) | 1.3 (1.1) | 0.7 (0.4) |
| UK (N=1,197) | 11.9 (4.6) | 4.5 (1.8) | 6.6 (2.1) | 3.9 (1.8) | 1.4 (1.1) | 0.7 (0.5) | 1.3 (1.2) | 0.7 (0.5) |
| GER (N=173) | 11.4 (3.8) | 4.3 (2.5) | 6.1 (2.7) | 3.8 (2.0) | 1.2 (0.7) | 0.5 (0.4) | 1.2 (1.1) | 0.5 (0.4) |
| FR (N=767) | 11.4 (4.6) | 4.1 (1.7) | 6.2 (2.3) | 3.6 (1.7) | 1.2 (1.0) | 0.5 (0.5) | 1.1 (1.2) | 0.5 (0.5) |
| HK (N=334) | 11.8 (4.2) | 4.7 (1.7) | 6.6 (2.3) | 4.2 (1.7) | 1.3 (0.8) | 0.6 (0.4) | 1.3 (1.0) | 0.6 (0.4) |
| SCP (N=823) | 10.8 (4.0) | 4.5 (0.8) | 6.2 (1.9) | 3.9 (1.1) | 1.0 (0.8) | 0.4 (0.3) | 1.1 (1.0) | 0.4 (0.3) |
| JP (N=1,292) | 12.3 (4.4) | 5.0 (1.5) | 7.0 (2.1) | 4.4 (1.3) | 1.3 (0.9) | 0.6 (0.4) | 1.3 (1.1) | 0.6 (0.4) |
| SG (N=657) | 12.0 (4.3) | 4.8 (1.5) | 6.9 (1.7) | 4.3 (1.5) | 1.4 (1.0) | 0.7 (0.5) | 1.4 (1.0) | 0.7 (0.5) |
| **Average (SD)** | **11.6 (4.4)** | **4.4 (1.6)** | **6.4 (2.1)** | **3.9 (1.5)** | **1.2 (0.8)** | **0.5 (0.4)** | **1.2 (1.1)** | **0.5 (0.4)** |

[a] Abbreviations: NY: New York State, CA: California, CAN: Canada, UK: United Kingdom, FR: France, HK: Hong Kong SAR, SCP: South China Provinces, JP: Japan, SG: Singapore

[b] Average amino acids mismatch against all available circulating strains in epidemic seasons from 2012/13 to 2018/19 in respective geographical region.

**Extended Data Table 12 | Prediction performance in retrospective data by geographical regions for HA of H3N2**

| Country/region[a] (sample size) | Average number of mismatch on HA sequence (out of 566 codons) (AAs) | | | | Average number of mismatch[b] on HA epitope (out of 131 sites) (AAs) | | | |
|---|---|---|---|---|---|---|---|---|
| | current-system (SD) | beth-1 (HA+NA) (SD) | LBI (HA) (SD) | beth-1 (HA) (SD) | current-system (SD) | beth-1 (HA+NA) (SD) | LBI (HA) (SD) | beth-1 (HA) (SD) |
| NY (N=1,287) | 11.1 (4.8) | 7.1 (3.5) | 8.3 (4.0) | 7.1 (3.5) | 7.3 (3.0) | 4.7 (2.3) | 5.4 (2.8) | 4.7 (2.3) |
| CA (N=1,011) | 10.5 (4.8) | 6.7 (3.5) | 8.2 (4.7) | 6.7 (3.5) | 6.9 (3.2) | 4.6 (2.5) | 5.3 (3.0) | 4.6 (2.5) |
| CAN (N=1,557) | 11.2 (4.0) | 8.6 (3.7) | 10.5 (4.9) | 8.6 (3.7) | 7.0 (2.6) | 5.5 (2.5) | 6.4 (3.4) | 5.5 (2.5) |
| UK (N=2,168) | 11.3 (5.2) | 7.6 (2.9) | 9.7 (5.1) | 7.6 (2.9) | 7.6 (3.3) | 5.3 (2.1) | 6.5 (3.4) | 5.3 (2.1) |
| GER (N=416) | 9.1 (2.8) | 7.5 (2.6) | 9.6 (4.5) | 7.5 (2.6) | 6.2 (2.1) | 5.1 (1.9) | 6.5 (3.0) | 5.1 (1.9) |
| FR (N=1,226) | 9.5 (2.8) | 7.0 (3.3) | 8.9 (4.8) | 7.0 (3.3) | 6.2 (2.4) | 4.9 (2.7) | 5.8 (3.3) | 4.9 (2.7) |
| HK (N=646) | 11.1 (4.6) | 7.9 (2.5) | 9.2 (4.6) | 7.9 (2.5) | 7.5 (3.1) | 5.7 (2.2) | 6.3 (3.2) | 5.7 (2.2) |
| SCP (N=1,687) | 12.2 (5.4) | 8.2 (2.3) | 10.4 (5.1) | 8.2 (2.3) | 8.2 (3.4) | 5.7 (1.9) | 6.7 (3.0) | 5.7 (1.9) |
| JP (N=2,394) | 11.6 (5.7) | 7.5 (2.9) | 9.6 (5.7) | 7.5 (2.9) | 7.5 (3.9) | 5.0 (2.3) | 6.2 (3.5) | 5.0 (2.3) |
| SG (N=1,040) | 12.0 (5.4) | 7.7 (2.7) | 9.8 (5.5) | 7.7 (2.7) | 8.0 (3.7) | 5.5 (2.0) | 6.5 (3.3) | 5.5 (2.0) |
| **Average (SD)** | **11.7 (5.1)** | **7.5 (2.2)** | **9.5 (4.7)** | **7.5 (2.2)** | **7.7 (3.4)** | **5.1 (1.7)** | **6.1 (2.9)** | **5.1 (1.7)** |

[a] Abbreviations: NY: New York State, CA: California, CAN: Canada, UK: United Kingdom, FR: France, HK: Hong Kong SAR, SCP: South China Provinces, JP: Japan, SG: Singapore

[b] Average amino acids mismatch against all available circulating strains in epidemic seasons from 2002/03 to 2018/19 in respective geographical region.

**Extended Data Table 13 | Prediction performance in retrospective data by geographical regions for NA of H3N2**

| Country/region[a] (sample size) | Average number of mismatch on NA sequence (out of 469 codons) (AAs) | | | | Average number of mismatch[b] on NA epitope (out of 12 sites) (AAs) | | | |
|---|---|---|---|---|---|---|---|---|
| | current-system (SD) | beth-1 (HA+NA) (SD) | LBI (NA) (SD) | beth-1 (NA) (SD) | current-system (SD) | beth-1 (HA+NA) (SD) | LBI (NA) (SD) | beth-1 (NA) (SD) |
| NY (N=1,129) | 7.3 (2.6) | 6.4 (4.4) | 6.6 (3.4) | 5.9 (3.8) | 1.0 (0.8) | 0.6 (0.7) | 0.7 (0.7) | 0.6 (0.7) |
| CA (N=904) | 7.1 (2.7) | 5.4 (3.5) | 6.6 (3.9) | 4.9 (2.5) | 0.9 (0.8) | 0.5 (0.7) | 0.6 (0.8) | 0.5 (0.7) |
| CAN (N=416) | 7.4 (2.4) | 5.6 (1.8) | 6.2 (3.3) | 5.2 (1.7) | 0.9 (0.9) | 0.5 (0.6) | 0.5 (0.6) | 0.5 (0.6) |
| UK (N=2,068) | 7.5 (2.5) | 5.9 (4.0) | 5.0 (1.6) | 5.5 (4.1) | 0.9 (0.8) | 0.5 (0.5) | 0.5 (0.5) | 0.5 (0.5) |
| GER (N=400) | 7.3 (2.7) | 5.8 (3.7) | 5.9 (2.8) | 4.9 (2.4) | 1.0 (0.8) | 0.5 (0.6) | 0.6 (0.7) | 0.4 (0.6) |
| FR (N=1,171) | 7.0 (2.7) | 5.2 (3.5) | 5.6 (2.3) | 4.5 (2.4) | 0.9 (0.8) | 0.5 (0.5) | 0.6 (0.6) | 0.4 (0.5) |
| HK (N=582) | 7.7 (2.8) | 5.5 (3.2) | 7.2 (3.6) | 5.1 (2.9) | 1.2 (0.9) | 0.6 (0.5) | 0.8 (0.7) | 0.5 (0.5) |
| SCP (N=1,475) | 8.1 (3.0) | 6.0 (3.8) | 8.2 (4.0) | 5.5 (3.5) | 1.2 (1.0) | 0.7 (0.7) | 0.9 (0.8) | 0.7 (0.7) |
| JP (N=2,084) | 7.9 (2.9) | 6.0 (3.8) | 7.6 (4.7) | 5.5 (3.1) | 1.0 (0.9) | 0.6 (0.6) | 0.7 (0.7) | 0.5 (0.6) |
| SG (N=1,032) | 8.2 (2.9) | 5.3 (3.5) | 7.1 (3.9) | 4.8 (2.7) | 1.1 (0.9) | 0.6 (0.5) | 0.7 (0.7) | 0.5 (0.5) |
| **Average (SD)** | **7.7 (2.5)** | **5.7 (3.0)** | **7.3 (3.6)** | **5.3 (2.3)** | **1.0 (0.8)** | **0.6 (0.5)** | **0.7 (0.6)** | **0.5 (0.5)** |

[a] Abbreviations: NY: New York State, CA: California, CAN: Canada, UK: United Kingdom, FR: France, HK: Hong Kong SAR, SCP: South China Provinces, JP: Japan, SG: Singapore

[b] Average amino acids mismatch against all available circulating strains in epidemic seasons from 2002/03 to 2018/19 in respective geographical region.

### Extended Data Table 14 | Prediction performance for the 2019/20 season

| Gene | Prediction time | Data used up to | Predicted strain | Evaluation sample size | # mismatch on full gene (SD) | | | | # mismatch on epitopes (SD) | | | |
|---|---|---|---|---|---|---|---|---|---|---|---|---|
| | | | | | current-system | beth-1 (two-protein) | answer strain | *p*-value | current-system | beth-1 (two-protein) | answer strain | *p*-value[a] |
| pH1N1 HA | 2019-10 | 2019-06 | A/Hong Kong/2655/2019 | 1,566 | 11.3 (0.6) | 4.7 (0.9) | 4.8 (0.7) | <0.001 | 2.9 (0.2) | 2.0 (0.4) | 2.1 (0.3) | <0.001 |
| | 2022-03 | 2019-03 | A/Hong Kong/98/2019 | | | 4.8 (0.7) | | <0.001 | | 2.1 (0.3) | | <0.001 |
| pH1N1 NA | 2019-10 | 2019-06 | A/Hong Kong/2655/2019 | 1,193 | 8.1 (0.3) | 2.6 (0.6) | 2.7 (0.5) | <0.001 | 1.0 (0.1) | 0.1 (0.1) | 0.1 (0.1) | <0.001 |
| | 2022-03 | 2019-03 | A/Hong Kong/98/2019 | | | 2.7 (0.5) | | <0.001 | | 0.1 (0.1) | | <0.001 |
| H3N2 HA | 2019-10 | 2019-06 | A/Hong Kong/2678/2019 | 1,875 | 16.1 (4.3) | 11.0 (3.9) | 9.2 (3.8) | 0.016 | 10.4 (3.1) | 7.1 (2.2) | 6.1 (2.4) | 0.029 |
| | 2022-03 | 2019-03 | A/Hong Kong/2926/2018 | | | 12.2 (3.4) | | 0.062 | | 8.2 (1.7) | | 0.191 |
| H3N2 NA | 2019-10 | 2019-06 | A/Hong Kong/2678/2019 | 1,671 | 9.3 (2.1) | 5.5 (2.4) | 5.5 (2.1) | 0.001 | 1.8 (0.6) | 0.6 (0.4) | 0.6 (0.4) | <0.001 |
| | 2022-03 | 2019-03 | A/Hong Kong/2926/2018 | | | 4.7 (2.3) | | <0.001 | | 0.6 (0.4) | | <0.001 |

[a] *t*-test on log mismatch between the current-system and beth-1 predictions.

**Extended Data Table 15 | Prediction performance by prospective validation in 2020/21 - 2022/23**

| Gene | Target season | Prediction time | Data used up to | Predicted strain | Evaluation sample size | # mismatch on full gene (SD) | | | | # mismatch on epitopes (SD) | | | |
|---|---|---|---|---|---|---|---|---|---|---|---|---|---|
| | | | | | | current-system | beth-1 (two-protein) | answer strain | $p$-value | current-system | beth-1 (two-protein) | answer strain | $p$-value[a] |
| pH1N1 HA | 2020/21 | 2020-02 | 2019-12 | A/Hong Kong/4380/2019 | 122 | 11.8 (4.5) | 8.4 (2.5) | 4.9 (6.0) | <0.001 | 3.8 (1.2) | 2.7 (1.1) | 1.3 (2.0) | <0.001 |
| | 2021/22 | 2021-02 | 2021-01 | A/NAGASAKI/8/2020 | 1,358 | 10.1 (2.8) | 10.2 (7.2) | 8.2 (8.1) | 0.545 | 3.6 (1.3) | 3.3 (2.5) | 2.3 (2.5) | <0.001 |
| | 2022/23 | 2022-02 | 2022-01 | A/Bangladesh/3210910005/2021 | 1,949 | 10.7 (1.9) | 4.1 (1.8) | 4.1 (1.9) | <0.001 | 3.0 (0.7) | 1.0 (0.7) | 1.0 (0.7) | <0.001 |
| pH1N1 NA | 2020/21 | 2020-02 | 2019-12 | A/Hong Kong/4380/2019 | 111 | 5.9 (2.0) | 2.8 (3.0) | 2.4 (3.6) | <0.001 | 0.2 (0.5) | 0.2 (0.5) | 0.2 (0.5) | 1 |
| | 2021/22 | 2021-02 | 2021-01 | A/NAGASAKI/8/2020 | 1,360 | 3.3 (2.1) | 8.3 (2.0) | 3.2 (2.8) | <0.001 | 0.2 (0.5) | 0.2 (0.5) | 0.2 (0.5) | 1 |
| | 2022/23 | 2022-02 | 2022-01 | A/Bangladesh/3210910005/2021 | 1,864 | 4.1 (0.4) | 2.5 (0.9) | 2.5 (1.5) | <0.001 | 0.2 (0.2) | 0.2 (0.2) | 0.2 (0.2) | 1 |
| H3N2 HA | 2020/21 | 2020-02 | 2019-12 | A/Hong Kong/4435/2019 | 493 | 19.3 (3.6) | 18.9 (4.3) | 5.7 (4.5) | 0.031 | 13.1 (2.6) | 13.6 (3.4) | 4.2 (3.3) | 0.553 |
| | 2021/22 | 2021-02 | 2021-01 | A/Bangladesh/0004/2020 | 15,476 | 13.1 (2.5) | 5.7 (2.3) | 4.2 (4.0) | <0.001 | 10.8 (1.9) | 4.2 (1.6) | 2.9 (2.7) | <0.001 |
| | 2022/23 | 2022-02 | 2022-01 | A/Arsal/5773/2021 | 5,592 | 9.8 (1.8) | 9.1 (2.1) | 7.7 (3.1) | 0.327 | 6.3 (1.5) | 5.9 (1.7) | 5.0 (2.2) | 0.485 |
| H3N2 NA | 2020/21 | 2020-02 | 2019-12 | A/Hong Kong/4435/2019 | 469 | 6.4 (1.4) | 5.4 (1.4) | 2.6 (1.8) | <0.001 | 1.2 (0.4) | 0.2 (0.4) | 1.2 (0.5) | <0.001 |
| | 2021/22 | 2021-02 | 2021-01 | A/Bangladesh/0004/2020 | 15,074 | 3.1 (1.5) | 2.3 (1.6) | 2.3 (2.0) | <0.001 | 1.3 (0.9) | 0.2 (0.5) | 0.2 (0.5) | <0.001 |
| | 2022/23 | 2022-02 | 2022-01 | A/Arsal/5773/2021 | 5,342 | 5.0 (0.9) | 3.9 (0.9) | 3.9 (1.0) | 0.025 | 0.4 (0.3) | 0.4 (0.3) | 0.4 (0.4) | 1 |

[a] *t*-test on log mismatch between the current-system and beth-1 predictions.

### Extended Data Table 16 | Summary table of fitted parameters

| | pH1N1 HA | | | | pH1N1 NA | | | | H3N2 HA | | | | H3N2 NA | | | |
|---|---|---|---|---|---|---|---|---|---|---|---|---|---|---|---|---|
| | HK[a] | SG | IN | TH | HK | SG | IN | TH | HK | SG | IN | TH | HK | SG | IN | TH |
| *p*-value of g-measure | 0.013* | 0.001** | 0.034* | 0.004** | 0.056 | <0.001*** | 0.025 | 0.006** | 0.087 | 0.002** | 0.023* | 0.002** | 0.012* | 0.001** | 0.050 | 0.160 |
| *p*-value of the F-statistic | 0.044* | 0.001** | 0.049* | 0.034* | 0.181 | <0.001*** | 0.065 | 0.046* | 0.196 | 0.001** | 0.077 | 0.010 | 0.042* | <0.001*** | 0.146 | 0.395 |
| multiple R-squared | 0.669 | 0.898 | 0.703 | 0.693 | 0.599 | 0.943 | 0.727 | 0.759 | 0.457 | 0.904 | 0.709 | 0.738 | 0.674 | 0.918 | 0.630 | 0.337 |
| region specific (θ, h)[b] | (0.7,0) | (0.8,2) | (0.7,4) | (0.9,0) | (0.6,6) | (0.5,6) | (0.6,0) | (0.2,8) | (0.8,5) | (0.9,2) | (0.6,7) | (0.5,2) | (0.9,6) | (0.5,7) | (0.3,8) | (0.5,4) |
| median transition time τ (month) | 21.8 (IQR 17.4-27.1) | | | | 13.3 (IQR 9.6-20.3) | | | | 17.4 (IQR 8.9-25.3) | | | | 11.8 (IQR 4.7-21.7) | | | |
| mean transition time τ (month) | 22.7 (SD 8.5) | | | | 16.0 (SD 10.3) | | | | 20.3 (SD 15.6) | | | | 14.6 (SD 12.0) | | | |

[a] HK: Hong Kong SAR, SG: Singapore; IN: India; TH: Thailand

[b] Region-specific $\theta$ is used to calculate transition time, and it is estimated by optimizing the fitness function of the g-measure and some covariates including mean temperature, absolute humidity and season against annual sero-positivity rate. The annual mean temperature (centigrade) and mean absolute humidity ($g/m^3$) were retrieved from Hong Kong Observatory via https://www.hko.gov.hk/contentc.htm for Hong Kong SAR, and Weather Underground via https://www.wunderground.com/ for Singapore, India and Thailand. The yearly number of positive cases and specimens were retrieved from Hong Kong Center for Health Protection (HKHCP) via https://www.chp.gov.hk/en/resources/29/304.html for Hong Kong SAR, and FluNet of Global Influenza Surveillance and Response System (GISRS) via https://www.who.int/tools/flunet for Singapore, India and Thailand. The annual sero-positivity rate is calculated as number of positive cases divided by number of specimens. These epidemiology data are provided in https://github.com/mwanglab/beth-1/tree/main/epidemic_data.

**Extended Data Table 17 | Detailed information of the reagents and cell lines used in this study.**

| Reagents or resources | Source | Identifier |
|---|---|---|
| Minimum essential medium (MEM) | Thermo Fisher Scientific | 31095029 |
| Penicillin-Streptomycin (PS) | Thermo Fisher Scientific | 15140122 |
| Trypsin-EDTA | Thermo Fisher Scientific | 25200056 |
| Fetal Bovine Serum (FBS) | Thermo Fisher Scientific | 26140079 |
| HEPES | Thermo Fisher Scientific | 15630080 |
| N-p-Tosyl-L-phenylalanine chloromethyl ketone (TPCK) | Sigma-Aldrich | 4370285 |
| HA and NA gene | Sangon Technology | N/A |
| PR8 8 plasmids | This study | N/A |
| phw2000 vector | [51] | N/A |
| TransIT®-LT1 Transfection Reagent | Mirus Bio | MIR 2306 |
| AddaVax™ | InvivoGen | vac-adx-10 |
| **Cell lines** | **Source** | **Identifier** |
| HEK 293T | N/A | N/A |
| Humanized MDCK cells | [49] | N/A |
